# Environment and HSP90 modulate MAPK stomatal developmental pathway

**DOI:** 10.1101/426684

**Authors:** Despina Samakovli, Tereza Tichá, Miroslav Ovečka, Ivan Luptovčiak, Veronika Zapletalová, Yuliya Krasylenko, George Komis, Olga Šamajová, Theoni Margaritopoulou, Loukia Roka, Dimitra Milioni, Polydefkis Hatzopoulos, Jozef Šamaj

## Abstract

Stomatal ontogenesis is a key element of plant adaptation aiming to control photosynthetic efficiency and water management under fluctuating environments ^1,2,3^. Development of stomata is guided by endogenous and environmental cues and is tightly coupled to overall plant growth ^1,2,3^. YODA signaling pathway is essential to stomatal lineage specification^4,5,6^ since it regulates the activities of transcription factors such as SPEECHLESS (SPCH)^7,8,9,10^. Heat-shock proteins 90 (HSP90s) are evolutionarily conserved molecular chaperones implicated in a broad range of signalling pathways being integrated in interaction networks with client proteins^11,12,13,14^. Herein, based on genetic, molecular, biochemical, and cell biological evidence we report that heat-stress conditions affect phosphorylation and deactivation of SPCH and modulate stomatal density. We show that genetic and physical interactions between HSP90s and YODA control stomatal patterning, distribution and morphology. We provide solid evidence that HSP90s play a major role in transducing the heat-stress response since they act upstream and downstream of YODA signalling, regulate the activity and nucleocytoplasmic distribution of MAPKs, and the activation of SPCH. Thus, HSPs control the stomatal development both under normal temperature and acute heat-stress conditions. Our results demonstrate that HSP90s couple stomatal formation and patterning to environmental cues providing an adaptive mechanism of heat-stress tolerance response and stomatal formation in Arabidopsis.

---

Elevated temperatures attributed to global climate changes affect plant development and have a great economic impact as they directly influence crop yield and productivity. Molecular mechanisms of plant response to elevated temperatures involve heat shock proteins (HSPs) that are essential for plant adaptation and survival. Plant thermotolerance depends on the timely expression and accumulation of HSPs^15,16^.

Stomata are small cellular pores in the plant epidermis mediating gas and water exchange through transpiration ^1,2,3^. Stomatal formation and distribution depend on environmental parameters and internal cues ^1,2,3^. SPCH is the major transcription factor (TF) controlling stomatal ontogenesis ^7,8,9,10^. An intracellular mitogen-activated protein kinase (MAPK) cascade including YODA (MAPKKK), MKK4/5 (MAPKKs) and MPK3/6 (MAPKs)^4,5,6^ is required for SPEECHLESS (SPCH) regulation and stomata formation which is also negatively regulated by brassinosteroids through BIN2 kinase that phosphorylates and inactivates YODA^17^.

Since the documented global warming concerns both increase of the average daily temperatures as well as increase of extreme temperatures observed during the day^18,19^ to simulate the latter we exposed young Arabidopsis seedlings (2 days post-germination) to 37^0^C for 2h each day for 7 days (Supplementary Fig. 1a) that represents a sub-lethal temperature not influencing seedling viability^20^. Interestingly, wild-type plants responded to repetitive heat stress treatments by decreasing the stomatal density while in *yda* mutants that overproduce stomata in clusters^4,5,6^, heat stress almost rescued the severe phenotype (Fig. 1a, c). HEAT SHOCK PROTEIN 90 (HSP90), an evolutionary conserved molecular chaperone, interacts with a large repertoire of signaling proteins including kinases and TFs^11,12,13,14^ and controls numerous biological processes including organismal development and stress responses ^11,12,13,14,21,22,23^.

**Figure 1.**
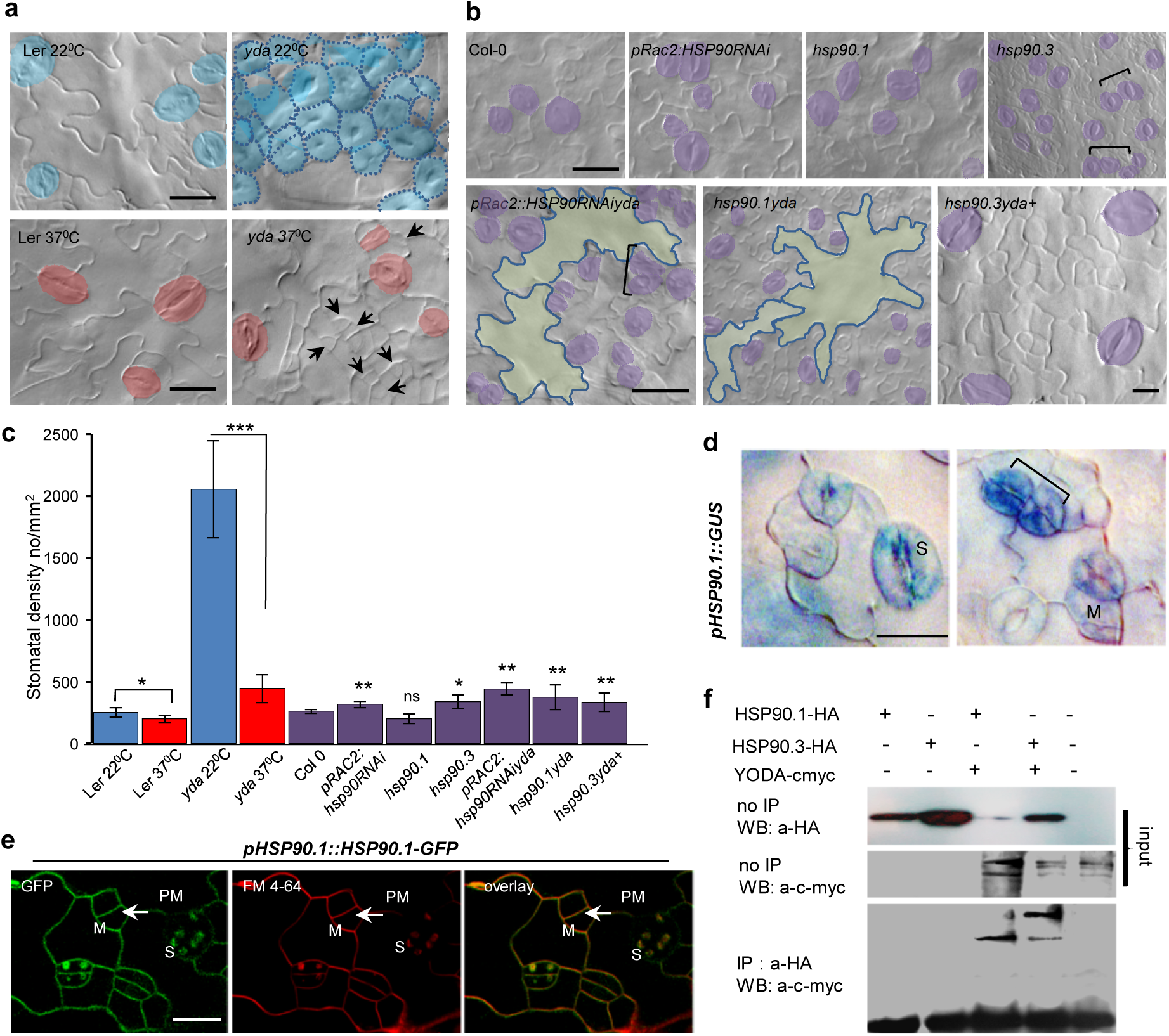
*HSP90s* are expressed in stomatal lineage cells and interact physically and genetically with YODA. **a** and **b**, Differential interference contrast (DIC) images of abaxial epidermis in 9 days post germination (dpg) cotyledons of indicated plant lines under control conditions or after heat stress. **a**, Col-0 and Ler represent wild types. **c**, Quantification of stomatal density in cotyledons of 9 dpg plant lines. Ten cotyledons of three independent biological repetitions were used each time. Brackets indicate clustered stomata and blue-marked pavement cells visualize big undivided cells. **d**, *pHSP90.1::GUS* expression in mature stomata and stomata precursors of 7 dpg cotyledons. **e**, Confocal microscope images of HSP90.1 localization using *pHSP90.1::HSP90.1-GFP* construct in cotyledons of 3 dpg seedlings showing predominant plasma membrane (PM) localization (arrows) in epidermal and stomatal lineage cells. S; stomata, M; meristemoids. **f**, Co-immunoprecipitation (IP) assays of the HSP90.1-HA and HSP90.3-HA with YODA-c-myc fusion protein. Protein extracts from *N. benthamiana* leaves co-transformed with the appropriate constructs were immunoprecipitated with anti-HA antibody and the immunoblots were probed with anti-c-myc antibody. The experiment was performed three times. Black arrows indicate meristemoids. Stomata and pavement cells in Figure 1a and 1b were artificially coloured for better visibility. Scale bars, 20 μm.

figTo rigorously test whether HSP90s were involved in the heat stress response of stomatal development we produced double mutants of *yda* with *hsp90.1*, a mutant in the heat stress-induced member, with *hsp90.3*, a mutant in almost constitutively expressed member, and with *pRAC2::HSP90RNAi* line (Fig. 1b and Supplementary Fig. 1b-d). Genetic impairment of *HSP90* genes and pharmacological inhibition of HSP90s with specific inhibitors, like GDA and 17-DMAG ^24,25^, caused stomatal clustering and higher stomatal density (Fig. 1b and Supplementary Fig. 2). Analysis of the stomatal densities in these double mutants revealed that *RNAi*-mediated depletion of HSP90s in the *yda* mutant attenuates stomata clustering in agreement with the pharmacological inhibition of HSP90s in the *yda* mutant (Fig. 1b, c, Supplementary Fig. 3a-c and Supplementary Fig. 4). Dramatic changes in cell division patterns, such as an increase of asymmetric cell divisions and a decrease of stomatal count, are characteristic for *pRAC2:HSP90RNAiyda* line (Supplementary Fig. 3a-c). Miss-patterned cell divisions and lower frequency of stomatal clustering were constantly observed in *hsp90yda* mutants (Fig. 1b, c, Supplementary Fig. 5, and Supplementary Fig 6). These-results reveal that the environmentally heat-induced proteins, HSP90s, act upstream of the YODA pathway and *HSP90* mutations are epistatic to *yda*.

**Figure 2.**
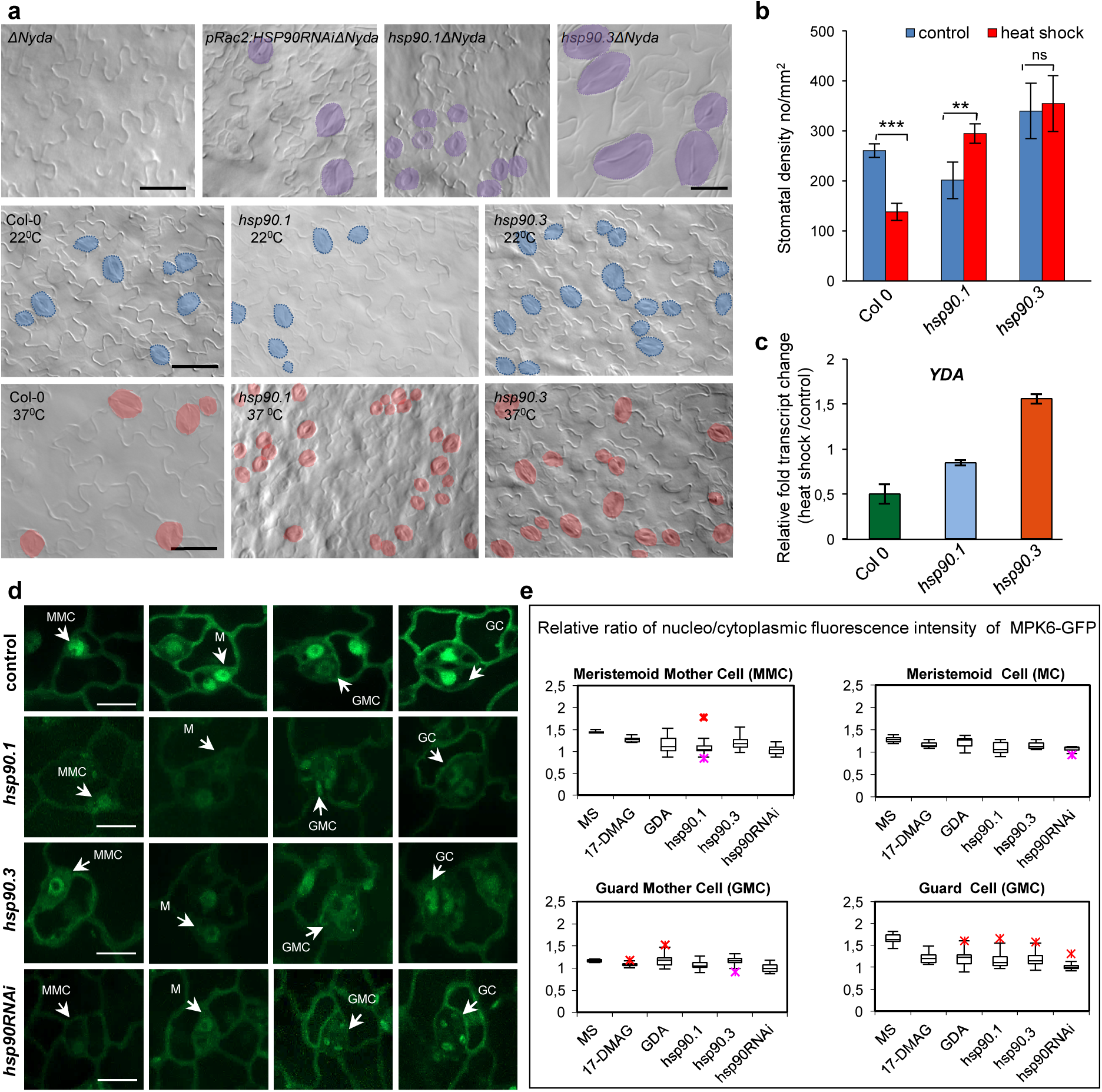
HSP90s act upstream of YODA and genetic depletions of HSP90s impact the subcellular distribution of MPK6 in the stomatal lineage cells. **a**, DIC images of abaxial epidermis in 9 dpg cotyledons of indicated plant lines grown under control conditions or after heat stress. **b**, Quantification of stomatal density in cotyledons of 9 dpg plant lines. Ten cotyledons of three independent biological repetitions were used each time. Brackets indicate clustered stomata and blue-marked pavement cells visualize big undivided cells. Data are means and error bars represent standard deviations *P<0.05, **P<0.01, n.s.-not statistically significant (one-tailed Student’s t-test). **c**, RT-qPCR analysis of *YDA* expression in heat stress-treated and non-treated wild-type (Col-0) and *hsp90.1* and *hsp90.3* mutants. **d**, *pMKP6::GFP-MPK6* localization in meristemoid mother cells (MMC), meristemoids (M), guard mother cells (GMC) and guard cells (GC) in the indicated genetic backgrounds. **e**, Quantification of the nucleo-cytoplasmic fluorescence intensity ratio of GFP-tagged MPK6 protein in control, *hsp90.1, hsp90.3* and *hsp90RNAi* genetic backgrounds. At least 15 cells per cell type were used for the fluorescence signal quantification in each case. Stomata cells in Figure 2a were artificially coloured for better visibility. Scale bars, 20μm for (a) and 10μm for (c).

To further address the role of HSP90s in the stomatal development, we showed that both *HSP90.1* and *HSP90.3* are expressed in stomatal cell lineage (SCL) from meristemoids to fully developed stomata (Fig. 1d, e, and Supplementary Fig. 7a, b) and verified that HSP90s physically interact with YODA kinase (Supplementary Fig. 7c-f).

The constitutive active YODA in *ΔNyda* mutants inhibits stomata formation (Fig. 2a)^4, 10^. Stomatal precursors were systematically formed in *ΔNyda* when HSP90s were pharmacologically inhibited partially rescuing the stomata-less *ΔNyda* phenotype, while fully developed stomata were occasionally encountered (Supplementary Fig. 4). *pRAC2:HSP90RNAi* based genetic depletion of HSP90s in *ΔNyda* exhibited few fully developed stomata, while stomatal precursors were abundant (Fig. 2a and Supplementary Fig. 3 a, b, d). Double mutants of *hsp90ΔNyda* showed stomatal complexes and SCL as in wild-type plants (Fig. 2a and Supplementary Fig. 8). Although, stomatal precursors were frequently observed in *hsp90ΔNyda*, as it was found in GDA-treated *ΔNyda* (Supplementary Fig. 4) stomata were sporadically detected having large guard cells (GCs) (Fig. 2a and Supplementary Fig. 8). The results show that HSP90s act also downstream of YODA in the signaling cascade for stomatal ontogenesis. The role of HSP90s in stomatal formation was further revealed as heat-stressed *hsp90.1* and *hsp90.3* mutants showed a counteracting impact on stomatal density (Fig. 2a, b). The counter effect can be explained by the significant induction of *HSP90.1* and *HSP90.3* transcripts especially in *hsp90.3* or *hsp90.1* mutants, respectively, caused by the repetitive cycles of heat stress (Supplementary Fig. 9). The existence of a feedback loop that enhances stomatal density in heat-shocked *hsp90.3* mutant was verified by the induced expression of *YDA* (Fig. 2c).

The transcriptionally-relevant function of MPK6 (e.g. through SPCH) during stomatal ontogenesis relies on its nuclear accumulation in SCL^26^. To further examine the role of HSP90s downstream of YODA, we tested how HSP90s affect the nucleocytoplasmic partitioning of MPK6 in all types of cells in SCL (Fig. 2d) ^27^. Both genetic and pharmacological depletion of HSP90s significantly decreased nuclear pools of MPK6 in MMCs, Ms and especially in GCs (Fig. 2d, e, and Supplementary Fig. 10) demonstrating that MPK6 translocation into the nucleus is affected by HSP90s. Predominant accumulation of MPK6 in the cytoplasm likely represses nuclear SPCH inactivation and leads to overproduction of stomatal precursors. In agreement, treatments with HSP90 inhibitors increased stomatal density and induced stomatal clusters in *mpk3-1 and mpk6-4* mutants (Fig. 3a, b).

**Figure 3.**
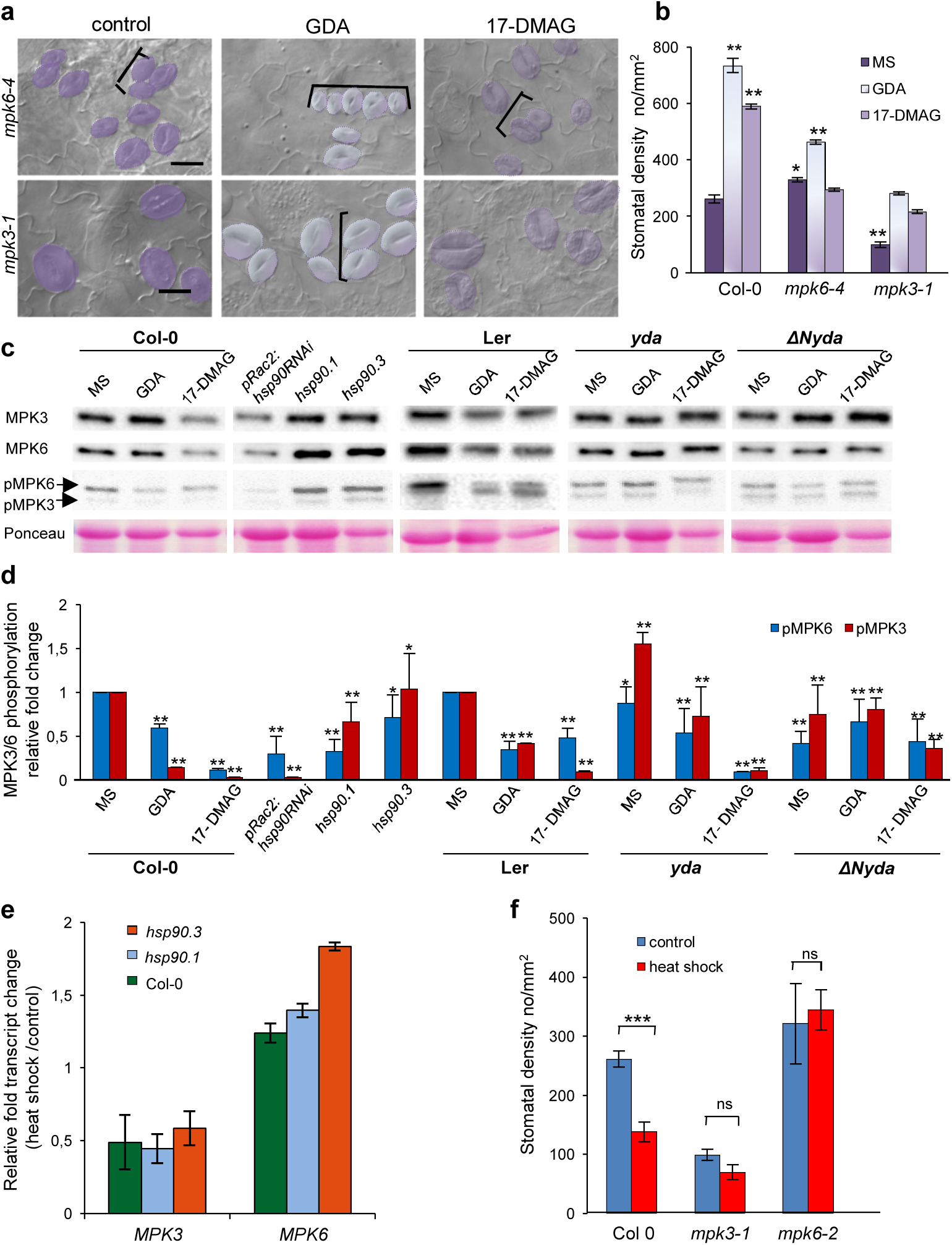
HSP90s depletion decreases the activation of MPK3/6 and induces stomatal formation. HSP90 pharmacological inhibition in *mpk6–4* and *mpk3–1*. **a**, DIC microscopy images of abaxial cotyledon epidermis of 9 dpg seedlings grown on plain MS medium or MS supplemented with GDA (5 μM), or 17-DMAG (5 μM). **b**, Stomatal density in 9 dpg wild type Col-0, *mpk6–4* and *mpk3–1* mutants under control conditions and after treatment with GDA and 17-DMAG (mean±S.D; *P<0.05, **P<0.01; one-way analysis of variance (ANOVA) followed by Tukey HSD test). Only statistical differences between the wild type plants grown under normal conditions against the rest are shown. **c**, Western blots of MPK3, MPK6 and phosphorylated MPK3/6 from protein extracts of 9 dpg plants. Full blots are available in Supplementary Fig. 12. **d**, Quantitative analysis of the MPK3/6 phosphorylation status in untreated and GDA or 17-DMAG-treated control and mutant lines. **e**, RT-qPCR analysis of *MPK3* and *MPK6* expression in heat stress-treated and non-treated wild-type (Col-0) and *hsp90.1* and *hsp90.3* mutants. **f**, Stomatal densities of heat shock treated and non-treated Col-0, *mpk3–1* and *mpk6–2* mutants. Data are means and error bars represent standard deviations ***P<0.001, (one-tailed Student’s t-test). Brackets indicate stomatal clusters. Stomata cells in Figure 3a were artificially coloured for better visibility. Scale bars, 50 μm.

The phosphorylation and concurrent activation of MPK3 and MPK6 within the YODA signaling pathway directly regulate downstream targets relevant to SCL specification, such as SPCH ^28,29^. Wild-type plants and *yda* mutant treated with HSP90 inhibitors as well as untreated *hsp90* mutants showed reduction of phosphorylated MPK3/6 pools (Fig. 3c, d). This result verified the role of HSP90s in the regulation of signaling components downstream of YODA. At the transcriptional level *MPK3* was downregulated, while *MPK6* was upregulated in wild type and *hsp90* mutants after heat-stress (Fig. 3e). Moreover, no changes in stomatal density were observed in *mpk3-1* and *mpk6-2* mutants after heat stress (Fig. 3f). These data suggest a critical interplay between MPK3/6 kinases and environment (heat stress) in the control of stomatal development.

In agreement with the stomatal phenotypic analysis, pharmacological depletion of HSP90s in the *yda* mutant decreased SPCH protein levels, while in the *ΔNyda* it resulted in an increase of SPCH protein abundance (Fig. 4a, b). Further, heat-stress treatment of wild-type plants reduced the SPCH protein abundance, while it showed no statistically significant impact on SPCH protein levels in the *pRAC2:HSP90RNAi, hsp90.1* and *hsp90.3* mutants (Fig. 4c, d), thus corroborating results of the phenotypical analyses (Fig. 2a, b). To further elucidate the molecular mechanism underlying heat-stress response we tested the protein levels of MPK3, MPK6, HSP90, SPCH and the activation of MAPKs. Immunoblot analysis revealed that the phosphorylation of MPK3 slightly increased (Fig. 4e, f), while MPK6 was strongly activated (15-fold increase) within 20 min of heat stress (Fig. 4e, f). Importantly, the pattern of MPK6 phosphorylation was also accompanied by increased SPCH phosphorylation showing a nearly 12-fold increase of phosphorylated to non-phosphorylated SPCH ratio (Fig. 4e, f). This suggests that activation of MPK6 is tightly coupled to SPCH phosphorylation during heat stress. Phosphatase treatments showed that the differential mobility of SPCH protein observed on immunoblots was due to the phosphorylation (Supplementary Fig. 12a). Next, we performed qPCR analysis for the major transcription factors controlling stomatal ontogenesis ^8^. Our results showed a moderate increase of *SPCH* transcript level in the *hsp90.1* mutant, while *MUTE* expression was almost unaffected and *FAMA* transcript level was upregulated under heat-stress conditions (Fig. 4g), suggesting an active interplay between acute heat-stress conditions and HSP90s. Taken together these results clearly show that HSP90s and MAPKs have a prominent role in the suppression of SPCH transcription factor that leads to the repression of stomatal development.

**Figure 4.**
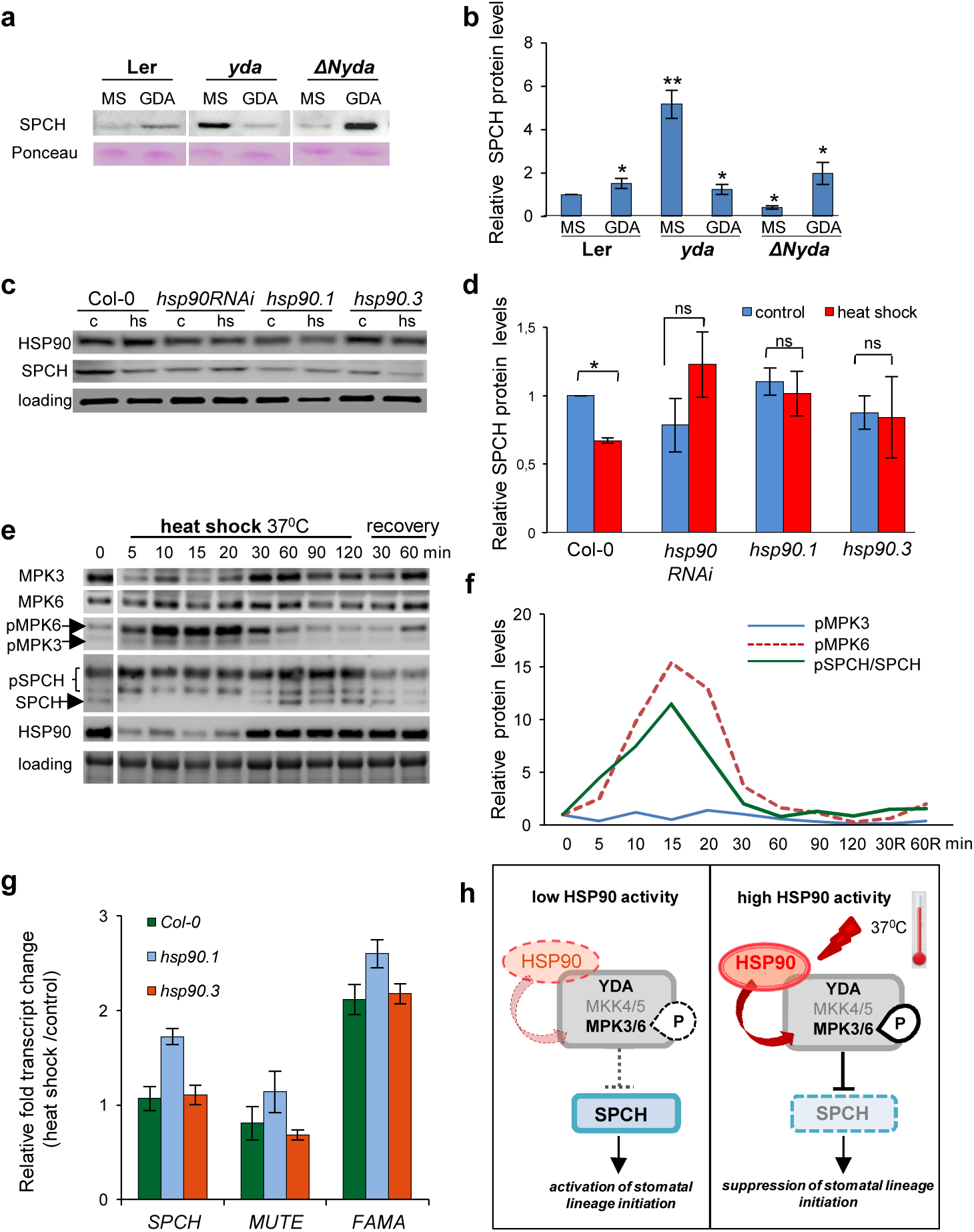
HSP90-dependent activation of MPK3/6 induces phosphorylation and destabilization of SPCH. **a**, Western blot of SPCH levels in designated 9 dpg seedlings grown in plain or 5 μM GDA-supplemented MS. **b**, SPCH protein levels in designated9 dpg seedlings grown in plain or 5 μM GDA-supplemented MS. **c**, Western blot analysis of SPCH and HSP90 in 9 dpg heat stress-treated and non-treated designated seedlings. **d**, Quantification of SPCH levels in heat stress-treated and non-treated plants **e**, Time course Western blot analysis of MPK3, MPK6, SPCH, HSP90 protein and phosphorylated MPK3/6. **f**, Relative phosphorylated MPK3/6 levels in the second day of heat shock. Relative fold changes of MPK3/6 protein levels were included in the calculations. **g**, qPCR analysis of *SPCH, MUTE* and *FAMA* in heat shock-treated and non-treated Col-0, *hsp90.1* and *hsp90.3* mutants. Data are means and error bars represent standard deviations *P<0.05, **P<0.01, n.s. - not statistically significant (one-tailed Student’s t-test). **h**, Proposed model for the modulation of YDA signaling cascade under depletion of HSP90 proteins and heat stress conditions that induce HSP90 activity. HSP90 integrate environmental stimuli and by regulating YDA cascade control the abundance and activity of SPCH, the master transcription factor fine-tuning leaf epidermis development.

This study reveals an intertwine relationship of acute environmental heat-stress conditions and HSP90s to regulate YODA signaling pathway during stomatal ontogenesis. We show that HSP90s act as upstream and downstream modulators of YODA, while MPK6 and SPCH phosphorylation is controlled by heat stress. Genetic and pharmacological inhibition of HSP90s decreased the phosphorylation of MAPKs and enhanced stomatal density under normal conditions. On the other hand, induction of HSP90s by heat stress suppresses stomatal formation due to the over-phosphorylation of MAPKs and SPCH (Fig. 4h). Considering the multi-parametric and complicated nature of heat stress response as a physiological process the contribution of additional signaling pathways cannot be excluded ^30^. In this report we demonstrate that HSP90s are crucial molecular determinants regulating the process of early SCL specification controlled by YODA, MAPKs and SPCH under regular and heat stress conditions.

## REFERENCES

1 Hetherington, A.M. & Woodward, F.I. The role of stomata in sensing and driving environmental change. Nature 424, 901–908 (2003).

2 Zoulias, N. et al. Molecular control of stomatal development Biochem J. 475, 441–454 (2018).

3 Nadeau, J.A. & Sack, F.D. Stomatal Development in Arabidopsis. Rockville, MD: American Society of Plant Biologists (2002a).

4 Wang, H. et al. Stomatal Development and Patterning Are Regulated by Environmentally Responsive Mitogen-Activated Protein Kinases in Arabidopsis. Plant Cell 19, 63–73 (2007).

5 Bush, S.M. & Krysan, P.J. Mutational evidence that the Arabidopsis MAP kinase MPK6 is involved in anther, inflorescence, and embryo development. J. Exp. Bot. 58, 2181–2191 (2007).

6 Komis, G. et al.. Cell and developmental biology of plant mitogen-activated protein kinases Annu. Rev. Plant Biol. 69, in press, https://doi.org/10.1146/annurev-arplant-042817-040314 (2018).

7 Pillitteri, L.J. & Torii, K.U. Mechanisms of stomatal development. Annu. Rev. Plant Biol. 63, 591–614 (2012).

8 Gray, J. E. Plant Development: Three Steps for Stomata. Curr Biol. 17, R213–R215 (2007).

9 Pillitteri, L.J. et al. Termination of asymmetric cell division and differentiation of stomata. Nature 445, 501–505 (2007).

10 Shpak, E.D. et al. Stomatal patterning and differentiation by synergistic interactions of receptor kinases. Science 309, 290–293 (2005).

11 Taipale, M., Jarosz, D.F. & Lindquist, S. HSP90 at the hub of protein homeostasis: emerging mechanistic insights. Nat. Rev. Mol. Cell Biol. 11, 515–528 (2010).

12 Rutherford, S.L. & Lindquist, S. HSP90 as a capacitor for morphological evolution. Nature 396, 336–342 (1998).

13 Queitsch, C. Sangster, T.A. & Lindquist, S. HSP90 as a capacitor of phenotypic variation. Nature 417, 618–624 (2002).

14 Samakovli, D. et al HSP90 canalizes developmental perturbation. J. Exp. Bot. 58, 3513–3524 (2007).

15 Larkindale, J. & Knight, M. R. Protection against heat stress-induced oxidative damage in Arabidopsis involves calcium, abscisic acid, ethylene, and salicylic acid. Plant Physiol. 128, 682–695(2002).

16 Hua, J. From freezing to scorching, transcriptional responses to temperature variations in plants. Curr. Opin. Plant Biol. 12, 568–573 (2009).

17 Kim, T.W. et al Brassinosteroid regulates stomatal development by GSK3-mediated inhibition of a MAPK pathway. Nature 482, 419–422 (2012).

18 Lewisa, S.C & King, A.D. Evolution of mean, variance and extremes in 21st century temperatures. Weather Clim. Ext. 15, 1–10 (2017).

19 Alexander, L.V. Global observed long-term changes in temperature and precipitation extremes: A review of progress and limitations in IPCC assessments and beyond. Weather Clim. Ext. 11, 4–16 (2016).

20 Silva-Correia, J. et al. Phenotypic analysis of the Arabidopsis heat stress response during germination and early seedling development. Plant Methods 10, 7 (2014).

21 Margaritopoulou, T. et al. HSP90 canonical content organizes a molecular scaffold mechanism to progress flowering. Plant J. 87, 174–87 (2016).

22 Samakovli, D. et al Brassinosteroid nuclear signaling recruits HSP90 activity. New Phytol. 203, 743–757 (2014).

23 Shigeta, T. et al Heat shock protein 90 acts in brassinosteroid signaling through interaction with BES1/BZR1transcription factor. J Plant Physiol. 178, 69–73 (2015).

24 Porter, J.R. et al. Discovery and development of Hsp90 inhibitors: a promising pathway for cancer therapy. Curr. Opin. Chem. Biol. 14, 412–420 (2010).

25 Jarosz, D.F. et al. Protein homeostasis and the phenotypic manifestation of genetic diversity: principles and mechanisms. Annu. Rev. Genet. 44, 189–216 (2010).

26 Zhang, Y. et al. The BASL polarity protein controls a MAPK signaling feedback loop in asymmetric cell division. Dev Cell 33, 136–149 (2015).

27 Smékalová, V. et al. Involvement of YODA and mitogen activated protein kinase 6 in Arabidopsis post-embryogenic root development through auxin up-regulation and cell division plane orientation. New Phytol. 203, 1175–1193 (2014).

28 Hunt, L. & Gray, J.E. The signaling peptide EPF2 controls asymmetric cell divisions during stomatal development. Curr. Biol. 19, 864–869 (2009).

29 Nadeau, J.A. & Sack, F.D. Control of stomatal distribution on the Arabidopsis leaf surface. Science 296, 1697–1700 (2002b).

30 Lau, O.S. et al. Direct Control of SPEECHLESS by PIF4 in the High-Temperature Response of Stomatal Development. Curr Biol, in press, doi.org/10.1016/j.cub.2018.02.054 (2018).

